# Tunneling Nanotubes Between Bone Marrow Stromal Cells Support Transmitophagy and Resistance to Apoptosis in Myeloma

**DOI:** 10.1101/2024.10.08.617234

**Authors:** Antonio Giovanni Solimando, Francesco Di Palma, Vanessa Desantis, Angelo Vacca, Maria Svelto, Francesco Pisani

## Abstract

Here we show that under oxygen-glucose deprivation (OGD), multiple myeloma cells transfer dysfunctional mitochondria to bone marrow stromal cells (BMSCs) for degradation (transmitophagy). These mitochondria are then transferred between BMSCs via tunneling nanotubes contributing to myeloma survival in OGD.

---

Multiple myeloma (MM) flourishes within the bone marrow (BM), a metabolically unique microenvironment with pronounced spatial oxygen-glucose gradients^1^.

Hypoxia and glucose deprivation drive metabolic reprogramming in MM cells^2^, reducing mitochondrial activity^3,4^ while enhancing their reliance on glycolysis via the Warburg effect. This metabolic shift fuels rapid ATP production and lactate buildup, creating an acidic niche that supports cancer progression. Targeting oxidative phosphorylation and glycolysis pathways may overcome the disease evolution, inhibiting the tumor growth in critical ecosystems^5^.

Consistently, it is known that a cell can transfer functional or non-functional mitochondria to other cells, contributing to mitochondrial quality control, thereby sustaining cellular plasticity^6,7,8,9^. Intercellular mitochondrial transfer in MM, supporting malignant cell survival, was previously investigated in normoxic co-culture systems, where the transfer of healthy mitochondria from BM stromal cells (BMSCs) to MM cells was observed^10,11^.

However, intercellular mitochondrial dynamic occurring under nutrient starvation and hypoxic conditions mimicking MM microenvironment remain unexplored.

Here, we set up a new co-culture system mirroring the MM niche by co-culturing MM cells and BMSCs in hypoxic and glucose-serum-deprived environment (0.2% O_2_; OGD), identifying a new mechanism involved in the survival of MM cells.

We found that MM cells survival is sustained by the transcellular degradation of unfunctional mitochondria, or transmitophagy^12^, that are transferred from MM cells to BMSCs, along with an intercellular transfer of MM cells mitochondria in post-fission state between BMSCs via Tunneling Nanotubes (TNTs). Notably, when TNTs between BMSCs are destroyed, BMSCs fail to support MM survival in OGD, highlighting the pivotal role of homotypic TNTs between BMSCs for MM progression.

JJN3 MM human cell line and BMSCs were cultured under normoxic control conditions (Nx) or in OGD. Mitochondrial membrane potential (ΔΨ), apoptosis, and TNTs were analyzed (Figure 1A). The ΔΨ of JJN3 cells was higher compared to that measured for BMSCs, and under OGD conditions, the ΔΨ of JJN3 cells strongly decreased, while the ΔΨ of BMSCs remained unaffected (Figure 1B). To mimic the hypoxic MM microenvironment, JJN3 cells were co-cultured with BMSCs under both normoxic and OGD conditions, and apoptotic cells were measured. In OGD BMSCs were found to be healthy and able to rescue JJN3 cells from OGD-triggered apoptosis (Figure 1C). Nx and OGD co-cultures were stained for F-actin and specifically analyzed for TNTs^13,14^. It was found that OGD triggers homotypic TNT formation between BMSCs (Figure 1D), while heterotypic TNTs between JJN3 cells and BMSCs were extremely rare. These homotypic TNTs were observed to be detached from the substrate (Figure 1E), capable of transferring cargo between BMSCs, and were highly dynamic (Figure 1F).

**Figure 1:**
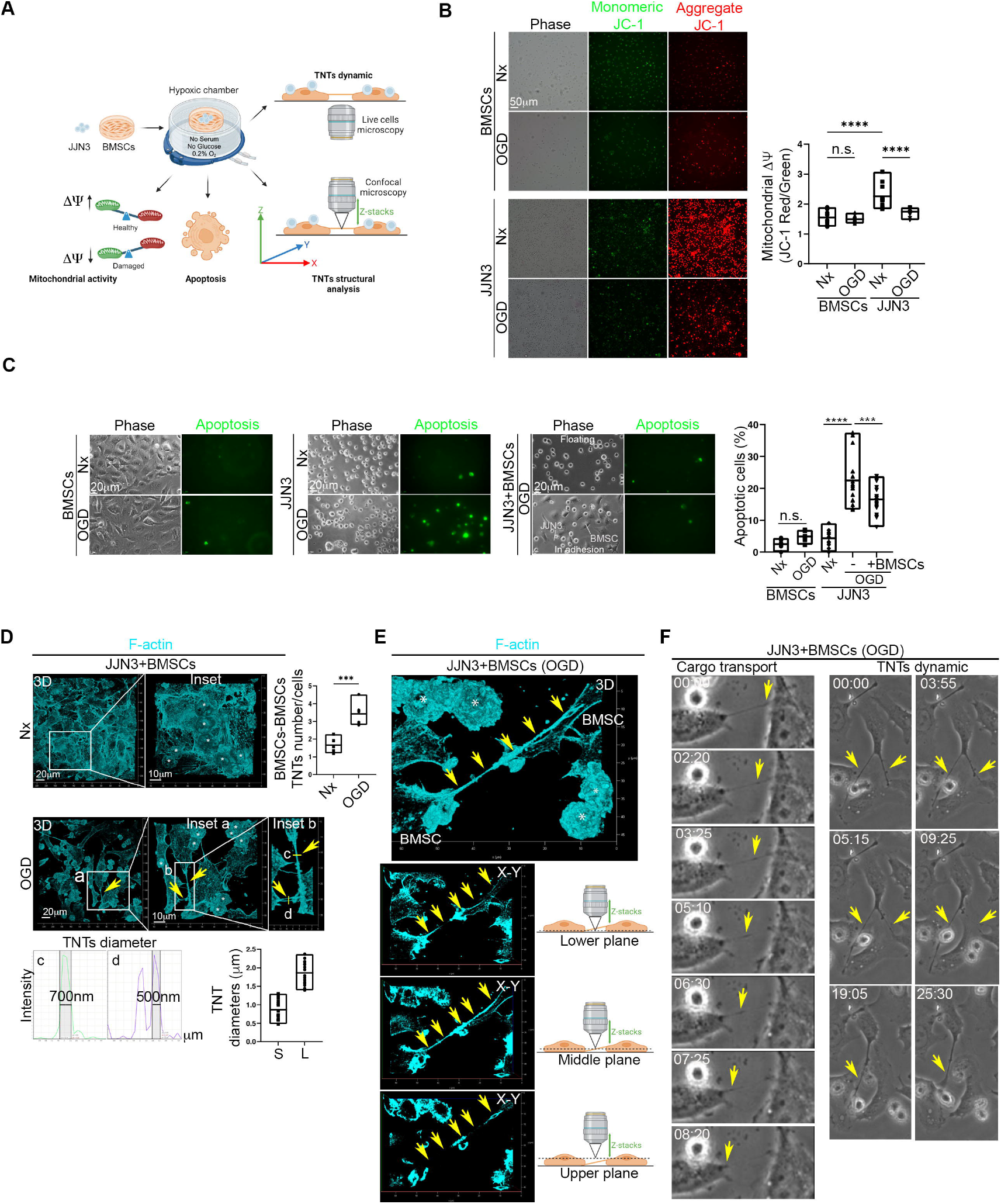
BMSCs Rescue JJN3 from OGD-Induced and Mitochondrial-Dependent Apoptosis and Generate an Extensive Homotypic TNTs Intercellular Network. **A**. Schematic representation of the Oxygen-Glucose Deprivation (OGD) co-culture model used here. JJN3 cells were co-cultured with BMSCs under control conditions or in the absence of serum, glucose, and in a hypoxic environment (0.2% O_2_). The co-culture was analyzed for mitochondrial membrane potential (ΔΨ), cell apoptosis, TNT structure (confocal microscopy), and TNT dynamics (live-cell microscopy). **B**. OGD strongly affects mitochondrial membrane potential (measured through JC-1; ΔΨ) in JJN3 cells, while BMSCs preserve ΔΨ under OGD conditions. **C**. JJN3 cells become apoptotic under OGD, while BMSCs are largely unaffected. When co-cultured with JJN3 cells, BMSCs rescue JJN3 cells from OGD-triggered apoptosis. **D-E-F**. OGD induces the formation of homotypic tunneling nanotubes (TNTs) between BMSCs cells. The diameters of small (S) and large (L) TNTs were measured (full width at half maximum, FWHM) (D). These TNTs were found to be detached from the surface (E) and transport cargo between BMSCs cells (F). In panels D and E, asterisks indicate JJN3 cells.

To trace the intercellular dynamics of JJN3 mitochondria, JJN3 cells were stained for mitochondria using the fixable Mitotracker Deep Red (JJN3 Mito) and co-cultured with BMSCs under Nx or OGD conditions. The cells were then analyzed by confocal microscopy to determine JJN3 mitochondrial localization (Figure 2A). Confocal microscopy revealed the presence of JJN3 mitochondria inside BMSCs. Under OGD conditions, the intercellular transfer of mitochondria increased, and the size of the transferred mitochondria was smaller compared to Nx, indicating that OGD triggers the transfer of mitochondria from JJN3 cells to BMSCs and that in OGD JJN3 mitochondria inside BMSCs were in a post-fission state (Figure 2B).

**Figure 2:**
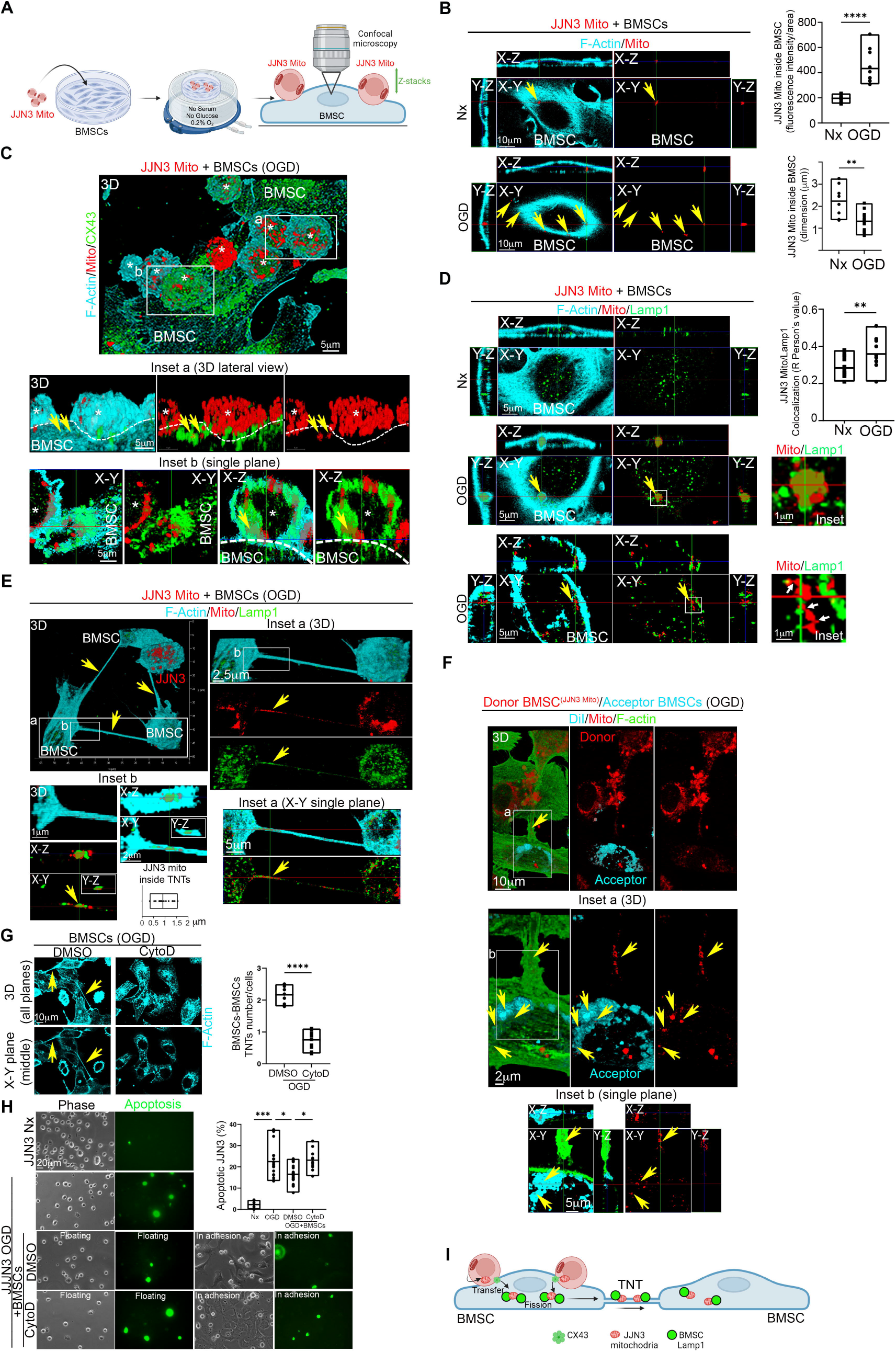
TNTs Between BMSCs Transfer JJN3 Mitochondria In Post-Fission State and Contribute to JJN3 Resistance to OGD. **A**. Schematic representation of the co-culture model used here. Mitochondria of JJN3 cells were stained with fixable Mitotracker Deep Red and co-cultured under normoxic (NX) or OGD conditions with BMSCs. The co-culture was analyzed for mitochondrial transfer and colocalization with Lamp-1 in BMSCs through confocal microscopy. **B**. JJN3 cells interact with BMSCs through CX43. Confocal microscopy of the OGD co-culture shows high CX43 abundance in the interaction region, and mitochondria frequently colocalize with CX43. **C**. Quantification of mitochondrial transfer from JJN3 cells to BMSCs under NX and OGD conditions. OGD triggers intercellular mitochondrial transfer for small post-fission mitochondria. **D**. OGD triggers the colocalization of JJN3 mitochondria and Lamp-1 in BMSCs. Note the presence of an actin cage typical of damaged mitochondria destined for mitophagy (PMID: 30058425) (central panel). Once inside BMSCs, multiple fission points were observed in JJN3 mitochondria (bottom panel, white arrows). **E**. TNTs between BMSCs in co-culture with JJN3 Mito under OGD conditions contain lysosomes and small JJN3 mitochondria in post-fission state. The co-culture was analyzed using high-resolution confocal microscopy for F-actin (cyan), JJN3 mitochondria (red), and Lamp-1 (green). Note the presence small mitochondria, with an average size of 1 ± 0.5 µm, originating from JJN3 cells inside TNTs between BMSCs. Lamp-1 frequently co-localizes with JJN3 mitochondria inside the TNTs. **F**. TNTs between BMSCs can transfer small JJN3 mitochondria in post-fission state from one BMSCs to another. A co-culture in OGD conditions, composed of donor BMSC (JJN3 Mito) (red) and acceptor BMSCs stained with DiI (Cyan), shows F-actin-positive (green) TNTs connecting donor and acceptor BMSCs (inset a) and the presence of JJN3 mitochondria inside the acceptor BMSCs (inset b). **G**. Cytochalasin-D (CytoD) in OGD destroys TNTs between BMSCs. **H**. TNTs between BMSCs contribute to JJN3 apoptotic resistance in OGD. Analysis of caspase 3/7 activity (green) reveals that pretreatment of BMSC with CytoD prevents the rescue of JJN3 cells in OGD conditions. **I**. Model of TNTs-mediated BMSCs-to-BMSCs JJN3 mitochondrial transfer supporting JJN3-to-BMSCs transmitophagy.

Analysis of gap junction protein Connexin 43 (Cx43) localization revealed that JJN3 cells were in direct contact with BMSCs via CX43 junctions, and JJN3 mitochondria appeared to be internalized by BMSCs through these CX43 junctions (Figure 2C), suggesting a CX43-dependent intercellular mitochondrial transfer^8^. Heterotypic TNTs between JJN3 cells and BMSCs were extremely rare, and no mitochondria were found inside them.

We further analyzed the co-culture for lysosomal-associated membrane protein 1 (Lamp1). Confocal microscopy clearly showed JJN3 mitochondria inside BMSCs interacting with Lamp1, with enhanced colocalization under OGD conditions. A more detailed analysis of JJN3 mitochondrial and Lamp1 localization and morphology in BMSCs revealed that, under OGD conditions, large masses of JJN3 mitochondria appeared to be internalized by BMSCs within an actin cage, surrounded by Lamp1, which is typical of damaged mitochondria destined for mitophagy^15^ (Figure 2D, central panel).

Once inside BMSCs, multiple mitochondria-Lamp1 contacts were observed, and fission points were identified in JJN3 mitochondria (Figure 2D, bottom panel, white arrows). Notably, lysosomal contacts mark the sites of mitochondrial fission^16^. Since JJN3 mitochondria lose membrane potential under OGD, a well-known trigger for mitophagy^17,18^, and various signs of fission appear in BMSCs for JJN3 mitochondria, we conclude that JJN3-to-BMSCs transmitophagy^12,15^ occurs under OGD conditions. We analyzed TNTs between BMSCs under OGD conditions for the presence of JJN3 mitochondria and Lamp1 localization. The analysis revealed that the homotypic TNTs network between BMSCs contains small JJN3 mitochondria in a post-fission state, along with Lamp1 (Figure 2E, inset a), and punctate JJN3 mitochondria interacting with Lamp1 (Figure 2E, inset b).

To assess whether TNTs between BMSCs can transfer these mitochondria between BMSCs, we co-cultured BMSCs with JJN3 mitochondria under OGD for 24 hours and washed out JJN3 cells, generating donor BMSCs^(JJN3 Mito)^. Subsequently, donor BMSCs^(JJN3 Mito)^ were incubated with acceptor BMSCs stained with DiI under OGD conditions. We found that donor BMSCs^(JJN3 Mito)^ were connected to acceptor BMSCs via TNTs containing JJN3 mitochondria in post-fission state and these mitochondria localize in acceptor BMSCs. These findings demonstrate that TNTs between BMSCs can transfer small JJN3 mitochondria in post-fission state between BMSCs (Figure 2F).

To test whether TNTs between BMSCs play a functional role in protecting JJN3 cells from OGD-triggered apoptosis, we disrupted the TNTs network between BMSCs in OGD using Cytochalasin-D (CytoD), a well-known drug that disrupts TNTs^9^. After washing out CytoD, we co-cultured CytoD pre-treated BMSCs with JJN3 cells under OGD conditions. Analysis of TNTs and cell apoptosis revealed that pre-treatment with CytoD destroy TNTs network between BMSCs (Figure 2G) and prevent the rescue of JJN3 cells from OGD-triggered apoptosis (Figure 2H). Taken together, our data show that the TNTs network between BMSCs allows the intercellular transfer of post-fission JJN3 mitochondria, supporting JJN3-to-BMSCs transmitophagy and contributing to the survival of JJN3 cells in the hypoxic MM niche (Figure 2I).

Our findings uncover a crucial metabolic adaptation in MM: we show that in a co-culture system mimicking the hypoxic MM milieu, the ΔΨ of MM cells is strongly affected and, most interestingly, these mitochondria are transferred to BMSCs for fission (transmitophagy). This mechanism has primarily been described in the central nervous system and has not been shown before in cancer^12,19,20^. Furthermore, in this context, BMSCs support MM cells survival by generating an active TNTs network among themselves for the transfer of MM mitochondria in a post-fission state, facilitating efficient transmitophagy within the BMSCs population. To the best of our knowledge, this is the first evidence of concerting role for transmitophagy and a TNTs network in the OGD MM microenvironment. Undeniably, treatments targeting oxidative phosphorylation, such as proteasome inhibitors are known to induce endoplasmic reticulum stress and apoptosis in MM cells^21^. Hence, resistance mechanisms driven by mitochondrial adaptations highlight the importance of metabolic plasticity. The quiescent MM cells, which are resistant to standard and novel agents, could correlate with metabolic shift sustaining the minimal residual disease in OGD^22,23^.

Overall, preventing TNTs-mediated intercellular mitochondrial exchange could weaken the protective environment that BMSCs provide to MM cells. This could represent a new frontier in MM therapy focused on metabolic vulnerabilities.

## Acknowledgments

This work was supported by by Unione Europea “National Center for Gene Therapy and Drugs based on RNA Technology”, PNRR missione 4 – componente 2 – investimento 1.4, cod.prog. CN00000041-CUP H93C22000430007” to F.P. and A.G.S and by the “Fondo per il Programma Nazionale di Ricerca e Progetti di Rilevante Interesse Nazionale—PRIN” (project n.2022ZKKWLW to A.G.S.). A.G.S. was also supported by a grant from “Società Italiana di Medicina Interna—SIMI” 2023 Research Award (CAMEL).

The graphical abstract and Figures 1A, 2A, and 2I were created with BioRender: BioRender.com/w13y615, Agreement number YH27EBHZVD.

## Authorship

Contribution: F.P., A.G.S., designed the research; F.D.P., V.D., F.P., and A.G.S. performed the experiments; all authors were involved in the analysis and interpretation of the data; A.G.S., F.P., F.D.P. and V.D. wrote the manuscript; F.P., A.V. and M.S. supervised the research and revised the manuscript. A.G.S. obtained the informed consent and obtained the primary samples. F.P. and A.G.S. financially supported the research. All authors approved the manuscript.

